# Impact of social dominance hierarchy on PACAP expression in the extended amygdala, corticosterone, and behavior in C57BL/6 male mice

**DOI:** 10.1101/2023.05.03.539254

**Authors:** Edward G. Meloni, William A. Carlezon, Vadim Y. Bolshakov

## Abstract

The natural alignment of animals into social dominance hierarchies produces adaptive, and potentially maladaptive, changes in the brain that influence health and behavior. Aggressive and submissive behaviors assumed by animals through dominance interactions engage stress-dependent neural and hormonal systems that have been shown to correspond with social rank. Here, we examined the impact of social dominance hierarchies established within cages of group-housed laboratory mice on expression of the stress peptide pituitary adenylate cyclase-activating polypeptide (PACAP) in areas of the extended amygdala comprising the bed nucleus of the stria terminalis (BNST) and central nucleus of the amygdala (CeA). We also quantified the impact of dominance rank on corticosterone (CORT), body weight, and behavior including rotorod and acoustic startle response. Weight-matched male C57BL/6 mice, group-housed (4/cage) starting at 3 weeks of age, were ranked as either most-dominant (Dominant), least-dominant (Submissive) or in-between rank (Intermediate) based on counts of aggressive and submissive encounters assessed at 12 weeks-old following a change in homecage conditions. We found that PACAP expression was significantly higher in the BNST, but not the CeA, of Submissive mice compared to the other two groups. CORT levels were lowest in Submissive mice and appeared to reflect a blunted response following social dominance interactions. Body weight, motor coordination, and acoustic startle were not significantly different between the groups. Together, these data reveal changes in specific neural/neuroendocrine systems that are predominant in animals of lowest social dominance rank, and implicate PACAP in brain adaptations that occur through the development of social dominance hierarchies.

## 1. Introduction

Stratification of social status has important implications for behavior and emotional health in both animals and humans, with individuals lower in social standing generally experiencing worse outcomes than those with higher standing^1–4^. Chronic psychosocial stress and agonistic interactions experienced by those in subordinate roles likely underlies this strong correlation between social rank and health^5, 6^. In laboratory mice, several different paradigms have been used to examine differences in neural systems and substrates that may correlate with the assignment of animals as either dominant (e.g. alpha animals) or subordinate (e.g. beta, gamma, delta animals) in social dominance ranking constructs^7–9^. These include behavioral assays such as the tube test, territory urine marking, and the warm spot test where induced conflict and ensuing dominance among animals determines rank order^10–12^. One of the most commonly used assays to identify social dominance rank is through the observation and scoring of agonistic behavior of group-housed animals in a homecage setting^10, 12–14^. This method relies on the natural, self-organizing dominance hierarchies that can develop in many strains of group-housed cages of mice as animals engage in periodic offensive (e.g. aggressive) and defensive (e.g. submissive) behaviors through the course of standard laboratory animal housing^14^. Agonistic behaviors can be elicited spontaneously, at the onset of the diurnal active period (e.g. lights-off), or after a simple perturbation such as a change into a new unfamiliar primary enclosure (e.g. cage change)^15^. Once established, hierarches are generally stable with especially high maintenance of rank over time for cagemates identified as either most-or least-dominant among group housed animals^7, 16, 17^. Using this methodology, several studies implicate medial prefrontal cortex-and nucleus accumbens-dependent neural circuits that influence social dominance rank^12, 18–20^. Further, alterations in neuropeptides, hormones, genes and inflammatory biomarkers that are differentially expressed in dominant versus subordinate cagemates have been identified^14, 15, 21–24^. Of interest to our own work studying stress peptides such as corticotropin releasing factor (CRF)^25–,27^, previous studies have shown alterations in CRF mRNA levels in dominant versus subordinate animals^14, 24^, suggesting a relationship between social dominance and this important regulator of stress and social behavior ^28–30^.

Along with CRF, another neuropeptide—pituitary adenylate cyclase-activating polypeptide (PACAP)—has been implicated in brain adaptations to stress^31–33^ and regulation of CRF and stress hormones such as corticosterone (CORT) through the hypothalamic-pituitary-adrenal (HPA) axis^34–36^. PACAP belongs to the secretin/glucagon superfamily of peptides and exists in two biologically active forms (as 38-and 27-amino acid peptides) found in peripheral tissues and brain^37^. PACAP-38 is the predominant form in the brain and shares identical amino acid sequence homology in species including mice, rats, sheep, and humans, indicating strong evolutionary conservation^38^. The high density of PACAPergic afferents and PACAP-type-I receptors (PAC1Rs) within the extended amygdala (e.g. central nucleus of the amygdala [CeA] and bed nucleus of the stria terminalis [BNST])^39–41^ suggests that this peptide plays a role in modulating neural activity related to psychological stress as well as experience-dependent learning^42–44^. We have demonstrated that PACAP influences AMPA receptor-dependent synaptic transmission in the CeA^45^, which receives direct PACAPergic innervation from the parabrachial nucleus^46^ and may where PACAP may be endogenously released in response to pain. Further, we found that exogenously administered PACAP can affect expression of fear-related behavior and CORT levels in a fear-conditioning paradigm^47, 48^ and impact motivation and social behavior^49^. The current study was designed to examine differences in PACAP expression within the extended amygdala in cagemates of mice ranked according to social dominance interactions. Further, we examined the impact of social dominance rank on acoustic startle response, a behavioral test that is sensitive to stress and emotional state and is influenced by PACAP^49, 50^. Given that preclinical paradigms have been useful tools to help understand the neurobiology of emotional disorders that may arise from repeated psychological or physical stressors, understanding how PACAP systems influence social dominance behavior may have face and construct validity for studying these illnesses as they appear in humans^4, 18, 51^.

## 2. Methods and Materials

### 2.1. Animals

Male C57BL/6 mice bred and housed in the McLean Hospital vivarium were used. To generate the experimental animals, timed breeding pairs were established to allow pooling of male offspring into group cages of 4 mice/cage at approximately 3 weeks of age. At the time of group housing, the body weight of each mouse was determined and four animals of similar weight were housed together in each polycarbonate cage (28 x 18.5 x 12.5 cm) with laboratory bedding (Alpha Chip; Northeastern Products Co.) and one square (5 x 5 cm) of nesting material (Nestlet; Ancare). All mice were maintained on 12/12 h light dark cycles (lights on at 700 h) and food and water were provided *ad libitum.* Weekly cage changes to place animals into a clean homecage with new bedding and nesting material occurred on the same day each week always between 1000 and 1300h. The timeline of all procedures is illustrated in **Figure 1A**. All animal procedures were approved by McLean Hospital’s Institutional Animal Care and Use Committee (Office of Laboratory Animal Welfare Assurance number A3685-01) in accordance with the National Institute of Health *Guide for the Care and Use of Laboratory Animals (8^th^ Edition)*.

**Figure 1.**
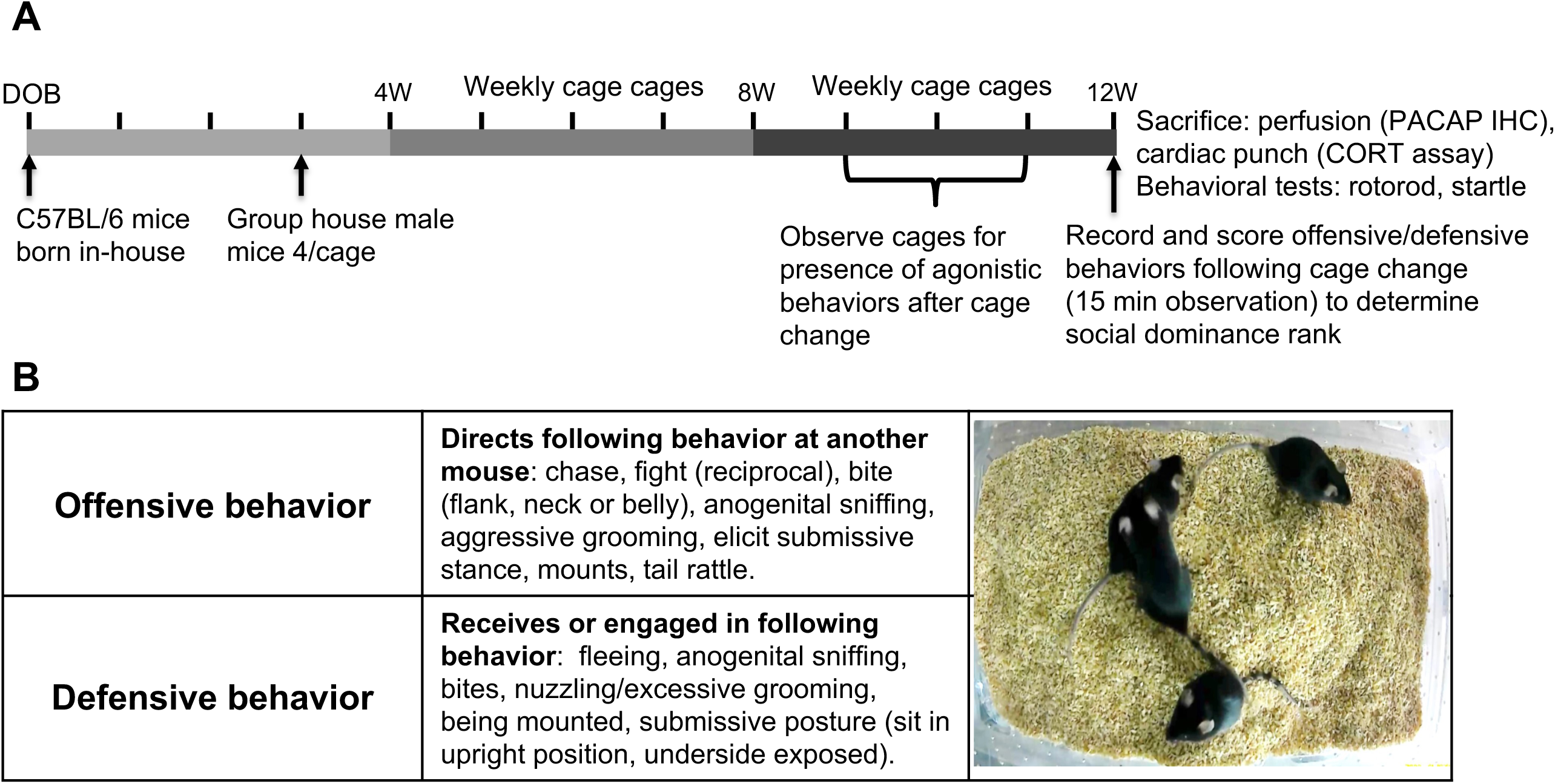
**(A)** Timeline of experimental design and procedures used in this study. **(B)** Definitions of agonistic behaviors used to rank social dominance based on number of observations of offensive and defensive behaviors within cages of group-housed mice (4/cage) following cage change at 12 weeks of age (W). Pituitary adenylate cyclase-activating polypeptide (PACAP); immunohistochemistry (IHC); corticosterone (CORT); date of birth (DOB).

### 2.2. Social interaction measurement and determination of social dominance hierarchy

Starting at 10 weeks of age, each the 4 group-housed mice were weighed and tails were marked with distinctive markings to individually identify each animal prior to cage change. Mice were observed for the presence of agonistic behaviors (see below) at this time point and cages where this behavior was not present were omitted from further study. At 12 weeks of age, immediately following the introduction of each group of mice into a new clean homecage, social interactions among the group-housed mice were video recorded for 15 min. A digital camera was placed above the center of the cage and used to record behavior for scoring of offensive (aggressive) and defensive (submissive) behavior using previously described methods to identify dominant and submissive mice^13, 14^. The types of agonistic behaviors used to rank animals according to an overall score in a matrix of offensive versus defensive behaviors is illustrated in **Figure 1B**. By assigning a “point” to each mouse in the cage for every offensive and defensive behavior it engaged in over the 15 min period, they could be ranked as Dominant or Submissive (e.g. having the highest score for offensive behaviors and lowest score for defensive behaviors or vice versa, respectively). As described in Horii et. al. (2017)^14^, we identified a third rank of mice as “Intermediate” describing mice with offensive/defensive scores in-between those identified as dominant and submissive. However, unlike the study of Horii et. al. (2017) ^14^ which excluded this group from analyses, we retained these intermediates for our analyses as a third comparator group comprising the two animals in the cage of four that were ranked neither as Dominant nor Submissive.

### 2.3. PACAP immunohistochemistry (IHC)

Mice were overdosed with sodium pentobarbital (130 mg/kg; IP) and upon loss of toe-pinch reflex they were perfused intracardially with 0.9% saline (25 ml) followed by 2% paraformaldehyde, 0.05% gluteraldehyde, and 0.2% picric acid in 0.1 M PBS (75 ml) pH7.4. The execution of successful perfusion and fixation, which is critical for having confidence in the reproducibility of IHC assays, was evaluated by confirmation of three visual observations: tail curl within 1 minute following introduction of the fixative solution, rigor in the head, legs and arms following perfusion, and the presence of yellow coloring (provided by the picric acid in the fixative) in the brain during removal. As a result, two mice in the Intermediate group were judged to have had poor perfusions and were removed from analysis. Brains were removed, stored for 3 d in a 30% sucrose/0.1 M PBS solution, and then cut serially in 30 µm coronal sections with every-other section from the BNST, and every-third section from the CeA placed in a 2-ml borosilicate glass vial (12 sections/vial for both brain areas) for processing of PACAP immunohistochemistry. All incubations were done on a rocker platform at room temperature.

Sections were incubated in 0.6% hydrogen peroxide in 0.1 M PBS for 30 min followed by 3 washes (5 min) in 0.1M PBS. Sections were preincubated in antibody medium (2% normal donkey serum, 1% bovine serum albumin, 00.3% Triton-X-100 in 0.1M PBS) for 2 h followed by incubation for 24 h with a rabbit polyclonal antibody against PACAP (1:1000; Penninsula Labs; #T-4473) diluted in antibody medium. The sections were washed with PBS and incubated for 1 h with a donkey anti-rabbit biotinylated secondary antibody (1:400; minimum species cross-reactivity; Jackson ImmunoResearch). The sections were washed with PBS and incubated for 30 min with the avidin-biotin complex (Vector Laboratories, Burlingame CA) and then incubated for 5 min in 3,3’-diaminobenzidine (DAB)/H_2_O_2_ (Sigma Fast; Sigma) as a chromogen for visualization of PACAP peptide through the BNST and CeA. Processed sections were mounted on microscope slides and coverslipped with Permount (Fisher Scientific, Pittsburgh, PA) and observed with a Zeiss Axioscope 2 (Zeiss, Oberkochen, Germany). Still frame images of three coronal sections from each brain were captured approximating BNST and CeA regions from the mouse brain corresponding to images 52, 53, and 54 (BNSTov) and 69, 70 and 71 (CeAL) from the Allen mouse brain atlas (mouse.brain-map.org)^95, 96^. As the CeAL has a much longer rostro-caudal extent than the BNSTov, we focused on caudal sections of the CeAL, which has denser PACAP expression than rostral sections (see **Supplemental Figure 1**). PACAP immunoreactivity in each brain area was quantified by calculating the optical density (O.D.) of pixels using ImageJ software for Macintosh (Scion Corp, Fredrick, MD, USA); ImageJ is a public domain, JAVA-based image processing program developed by the National Institutes of Health (NIH). Captured sections were switched to gray scale for threshold adjustments (scale 0-141 units normalized to background values of white matter) and mean values of O.D. were calculated from 0.1 mm^2^ regions with a fixed template placed within each of the brain areas (see **Figure 2A**). Some alternate sections through the BNST and CeA were also labelled for PACAP using immunofluorescence according to previously published methods^48^ for illustration purposes and to further delineate the boundaries of the BNSTov and CeAL (**Figure 2B & D**) but were not used for quantification of PACAP in these regions.

**Figure 2.**
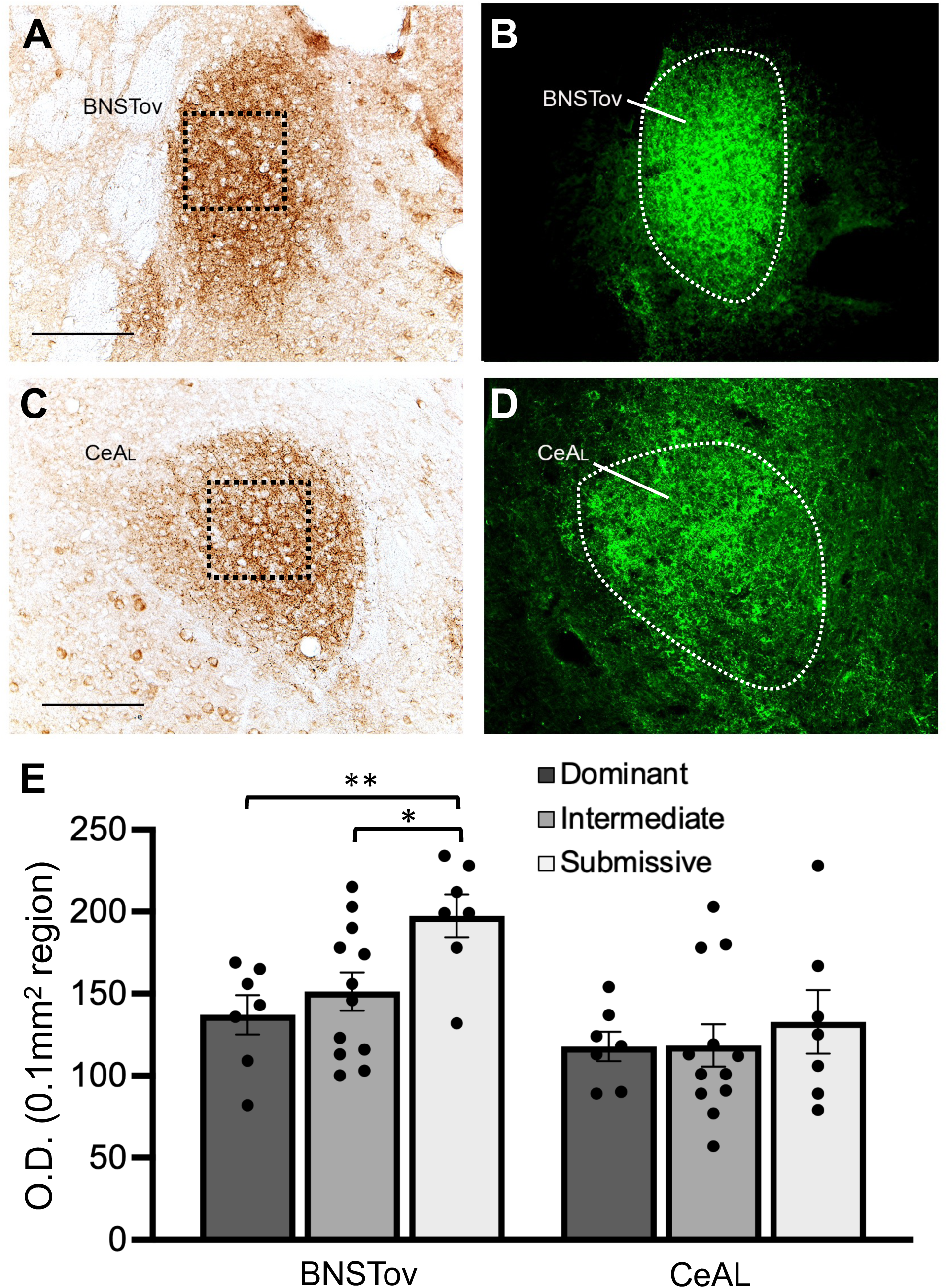
Representative coronal sections through the BNST (**A, B**) and CeA (**C, D**) showing PACAP expression; fluorescent images in **B** and **D** illustrate the approximate boundaries of the BNSTov and CeAL respectively. Optical density (O.D.) measurements of PACAP expression in 0.1 mm^2^ regions (boxes in A and C) in the BSNTov and CeAL were analyzed from brains of animals identified as Dominant, Intermediate or Submissive from scoring of social dominance interactions at 12 weeks of age. **(E)** Average O.D. values of PACAP expression in the BNSTov and CeL for each group. Animals in the Submissive group showed significantly higher levels PACAP expression in the BNSTov compared to the other two groups; PACAP expression was not significantly different in the CeAL between groups. Bar graph data are shown as mean±s.e.m. **P<0.005, *P<0.05. Scale bars = 100 μm.

### 2.4. Corticosterone assay

Serum CORT levels were measured following cage change and the 15 min session for recording of agonistic behavior in a separate cohort of group-housed 12 week-old mice to determine the impact of agonistic behavior on this stress-dependent hormone in animals of different social rank. Given diurnal variations in mouse CORT levels, blood sampling was conducted between 1000 and 1100h, when CORT levels are stably low^52, 53^. Mice were overdosed with sodium pentobarbital (130 mg/kg; IP) and upon loss of toe-pinch reflex, the chest cavity was opened. A 0.5 cc Insulin syringe (U-100 syringe; Becton-Dickson, Franklin Lakes, NJ) with a 28 gauge needle (0.5 in length) was used to draw blood from the right ventricle of the heart. This procedure is rapid (<3 min) and previous work indicates that it is unlikely that anesthesia significantly impacts CORT levels on this time scale^55^. Blood was transferred to a sterile 2 ml serum blood collection tube (BD Vacutainer; Becton-Dickson, Franklin Lakes, NJ) and allowed to clot at room temperature for 30 minutes before centrifugation for 10 min at 3000 rpm. Serum was removed, aliquoted, and stored at −80°C until assayed by ELISA following the manufacturer’s directions for quantitative determination of CORT levels in rat/mouse serum (Alpco Diagnostics, Salem NH). All samples were loaded onto a single plate in duplicate wells and assayed using a BioTek Synergy HT microplate reader to compare samples against a standard curve of known mouse CORT concentrations. The sensitivity of the assay was 6.1 ng/ml.

### 2.5. Rotorod

To explore whether motor capacities differ among groups, mice were tested on the accelerating rotorod (Ugo Basile; RotaRod model 7750) one day after the 12 week cage change and recording of social interaction among the group-housed mice. The four mice from each cage were placed on the rotorod cylinders at the same time starting at a slow rotational speed (4 rpm) which gradually increases over 2 min to a maximum of 40 rpm. Each lane on the device is equipped with individual timers to record latency-to-fall with a maximum trial length of 3 min. Mice are tested three times with a 5 min interval between tests with an overall latency-to-fall score averaged across the three trials for each mouse. Following this test, mice were perfused as described above for analysis of PACAP IHC.

### 2.6. Acoustic startle

A separated cohort of mice was tested for acoustic startle one day after the 12 week cage change and recording of social interaction among the group-housed mice. Testing was conducted in 4 identical mouse startle cages consisting of 6 x 6 x 5 cm Plexiglas cages with metal rod flooring attached to a load-cell platform. Both the startle cages and platform were located within a 69 x 36 x 42 cm fan-ventilated sound-attenuating chamber (Med Associates, Georgia, VT). Cage movement resulted in a displacement of a transducer in the platform where the resultant voltage was amplified and digitized on a scale of 0 to ±2000 arbitrary units by an analog-to-digital converter card interfaced to a personal computer (PC). Startle amplitude was proportional to the amount of cage movement and defined as the maximum peak-to-peak voltage that occurred during the first 200 ms after the onset of the startle stimulus. Constant wide-band background noise (60 dB; 10-20 kHz) and 50 ms startle stimuli (1-32 kHz white noise, 5 ms rise/decay) were generated by an audio stimulator (Med Associates) and delivered through speakers located 7 cm behind the startle cage. The calibration, presentation, and sequencing of all stimuli were under the control of the PC using specially designed software (Med Associates). For testing, mice were placed in the startle chambers and given a 5 min acclimation period followed by presentation of two habituating startle stimuli (100 dB, 30 s interstimulus interval; ISI). Mice were then presented with 30 startle stimuli at three different intensities (95, 100, 105 dB); the 10 trials at each intensity were presented in a semirandom order with a 30 s ISI. Startle amplitude data were expressed as the mean averaged across the 10 trials for each of the three startle-eliciting intensities.

### 2.7. Statistical Analyses

Data are presented as means ± standard error (SEM) for mice ranked as Dominant, Intermediate, or Submissive based on their agonistic behavior scores following the Week-12 cage change. The impact of social dominance rank on PACAP expression in the BNSTov and CeAL was analyzed using a two-way ANOVA with rank (Dominant, Intermediate, Submissive) as a between-subjects factor and brain area as a within-subjects factor. Body weight and rotorod performance was analyzed using separate independent-measures one-way ANOVAs. Startle data were analyzed using a two-way ANOVA with rank (Dominant, Intermediate, Submissive) as a between-subjects factor and startle intensity (95, 100, 105 dB) as a within-subjects factor. For analyses yielding significant main effects, subsequent post-hoc comparisons were made using Sheffe’s tests.

## 3. Results

### 3.1. PACAP IHC, body weight and rotorod performance

A total of seven cages of mice were scored for agonistic behaviors at 12 weeks of age to rank animals into Dominant, Intermediate and Submissive groups and process their brains for PACAP IHC in the extended amygdala. **Figure 2** illustrates representative light and fluorescent microscopy of brain sections through the BNST and CeA illustrating PACAP IHC in the BNSTov and CeAL subdivisions and quantification by optical density measurements of 0.1 mm^2^ regions in each of these areas for each social dominance group. A two-way ANOVA of PACAP expression across groups showed a main effect of brain area (*F*_1,23_ = 20.8, P < 0.0001), indicating generally higher levels of PACAP expression in the BNSTov versus the CeAL, and social dominance group (*F*_2,23_ = 3.53, P < 0.05). The brain area x social dominance group interaction was not significant. Subsequent post-hoc comparisons after significant main effects revealed significantly higher levels of PACAP expression between Submissive and Dominant (P<0.005) and Submissive and Intermediate (P<0.05) animals in the BNSTov but no pairwise differences in the CeAL between groups.

**Figure 3A** illustrates average body weight data from animals in the different social dominance groups analyzed for PACAP IHC in the BNSTov and CeA shown in **Figure 2**. A one-way ANOVA across groups revealed no significant main effect. Although there was a trend for Submissive animals to have lower average weight than Dominant or Intermediate animals, post-hoc comparisons with Sheffe’s test revealed no significant pairwise differences in body weight. We further analyzed these data to look for correlations between PACAP expression in the BNSTov – which was significantly higher in Submissive animals compared to the other two social dominance groups – and body weight; **Figure 3B** illustrates this relationship. There was a trend for animals with lower body weight to have higher BNSTov PACAP expression levels (r^2^=0.34), but this correlation was not significant. **Figure 3C** illustrates rotorod data from these same animals in the different social dominance groups. A one-way ANOVA across groups revealed no significant main effect of latency to fall, the dependent measure of motor coordination in this test.

**Figure 3.**
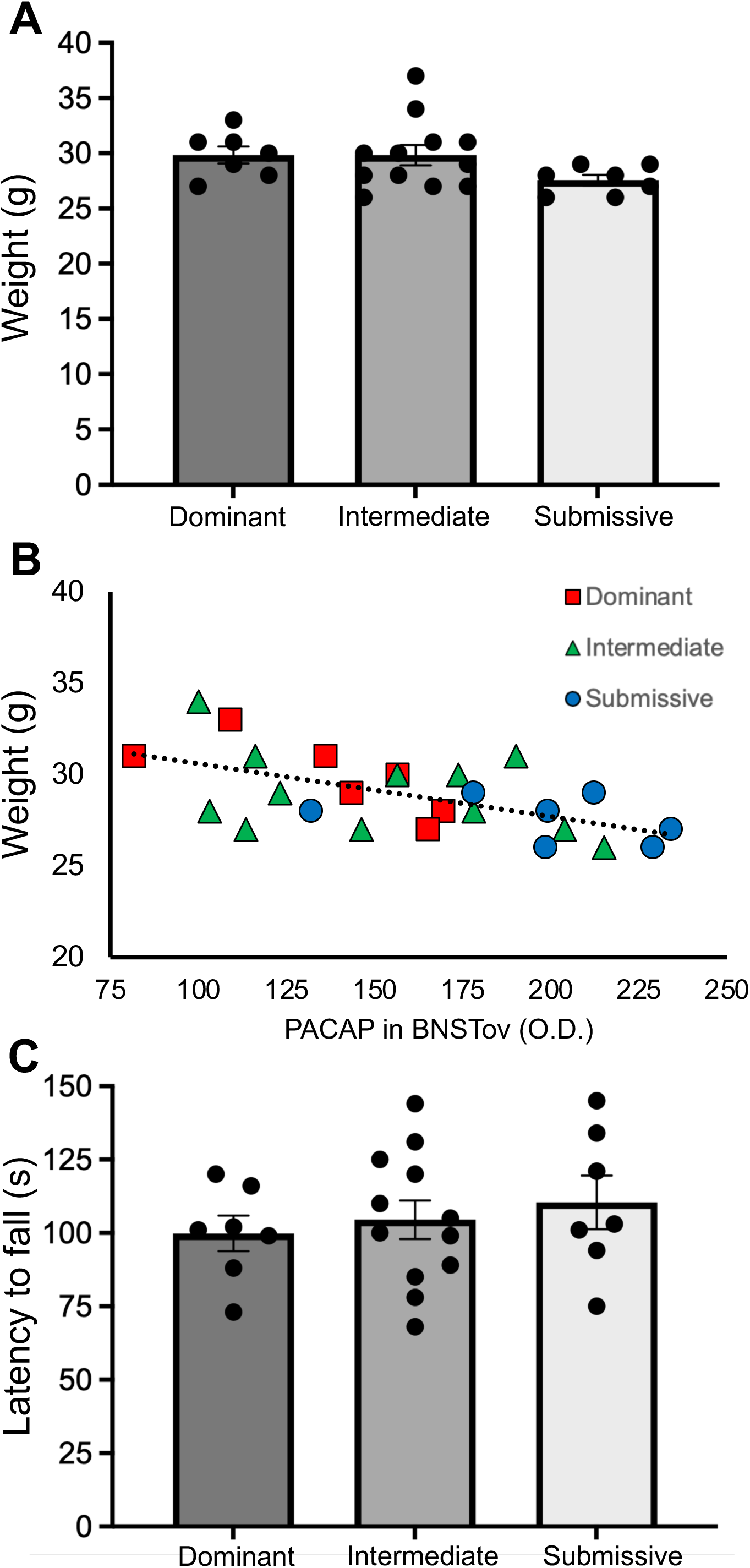
**(A)** Average body weight data from animals in the different social dominance groups at 12 weeks of age. **(B).** Relationship between body weight data and BNSTov PACAP expression (O.D.; optical density value from 0.1 mm^2^ region) for animals in the different social dominance groups; there was an overall trend for animals with lower body weight to have higher BNSTov PACAP expression levels (r^2^=0.34), but this correlation was not significant. **(C)** Average rotorod data from these same animals in the different social dominance groups. Bar graph data are shown as mean±s.e.m.

### 3.2. Serum CORT levels after agonistic interactions

A total of six cages of mice were used to examine blood serum CORT levels in Dominant, Intermediate and Submissive groups sacrificed immediately after animals engaged in agonistic interactions and behavior was recorded for identifying Dominant, Intermediate and Submissive animals following cage change at 12 weeks. CORT data for each of the social dominance groups are shown in **Figure 4**. A one-way ANOVA across groups revealed a significant main effect of CORT level (*F*_1,21_ = 8.29, P < 0.005). Subsequent post-hoc comparisons after significant main effects revealed significantly lower levels of CORT in Submissive animals compared to Dominant and Intermediate animals (P<0.005 both comparisons). An additional cage of 4 mice was sacrificed at 12 weeks, but prior to the weekly cage change, to provide a reference level for CORT in an unstimulated condition. Because we used the Week-12 cage change as the perturbation to stimulate agonistic behavior and score mice for social dominance, mice in this cohort were not individually identified as Dominant, Intermediate or Subordinate and CORT values were averaged to provide a reference level for basal CORT across all mice in the cage. The dashed line in **Figure 4** represents this reference value (92.8 ng/ml).

**Figure 4.**
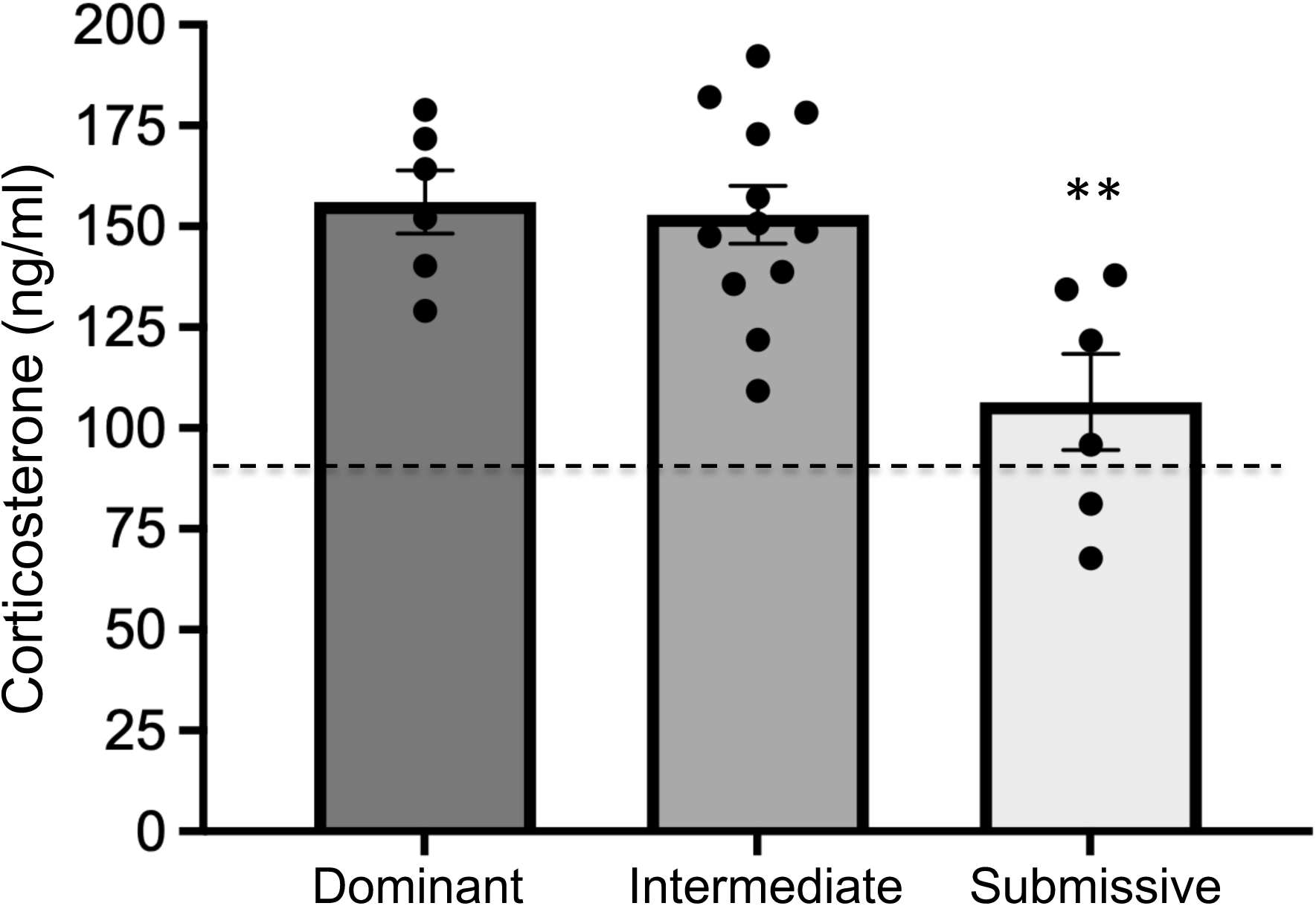
Serum corticosterone (CORT) levels from animals in the different social dominance groups measured at 12 weeks. Animals were sacrificed 15 min following the homecage change and recording session to identify Dominant, Intermediate and Submissive mice. Animals in the Submissive group showed significantly lower levels of CORT compared to other two groups (**P<0.005 versus both groups) following this session. Dashed line represents reference value (92.8 ng/ml) for basal CORT levels averaged across a separate cage of group-housed mice (4/cage) which did not receive cage change before being sacrificed. Bar graph data are shown as mean±s.e.m.

### 3.3. Social dominance and acoustic startle

**Figure 5** shows acoustic startle data from 7 cages of mice ranked as Dominant, Intermediate and Submissive. A two-way ANOVA of startle amplitude for each of the startle eliciting intensities (95, 100 105 dB) across groups showed only a main effect of startle intensity (*F*_2,50_ = 309.2, P < 0.0001) as startle amplitude is increased with more intense startle stimuli. Subsequent post-hoc comparisons did not reveal any significant differences in startle amplitude between social dominance groups at any startle intensity.

**Figure 5.**
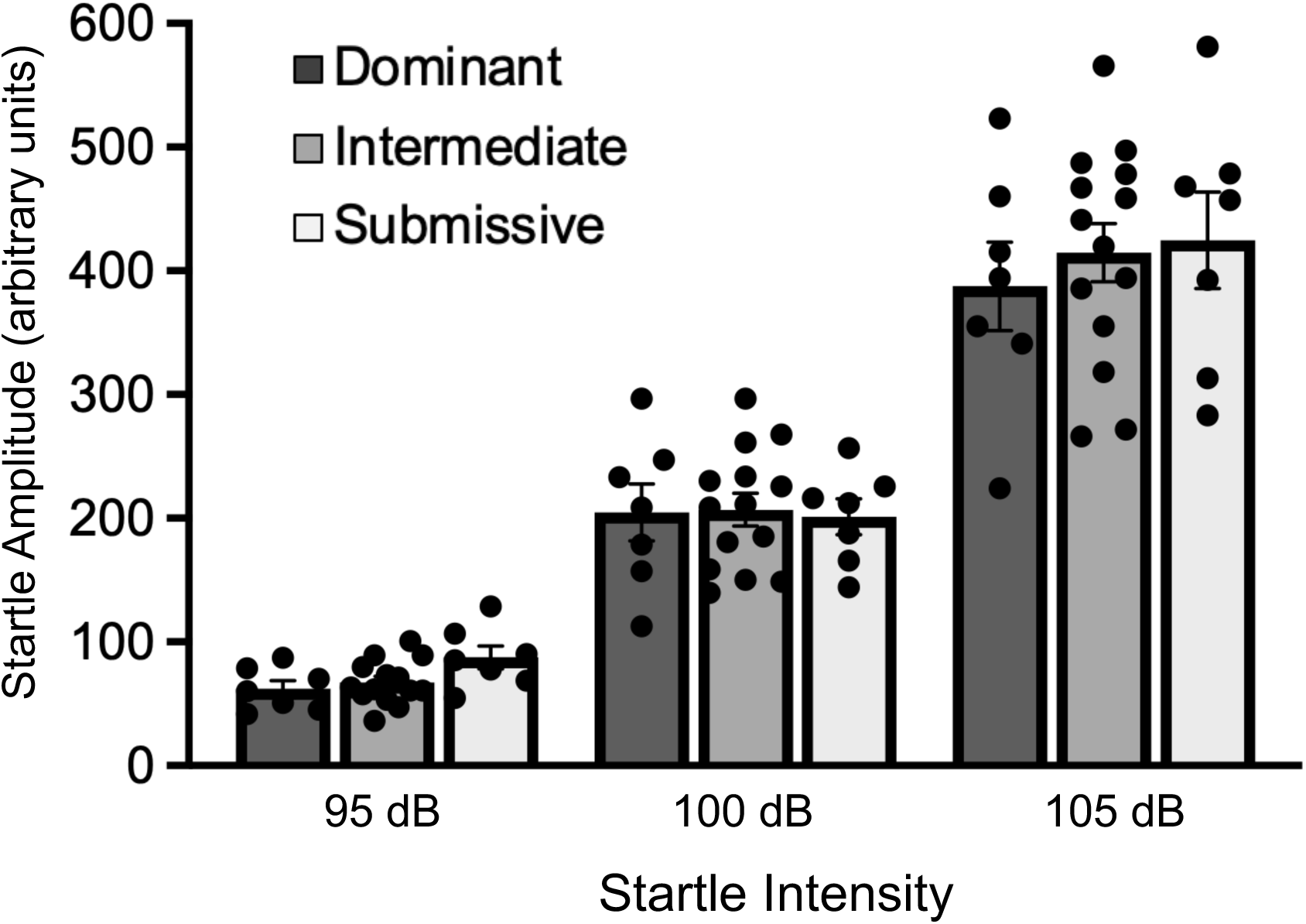
Acoustic startle response from animals in the different social dominance groups across startle intensity showing no differences between groups in this measure of sensorimotor reactivity. Bar graph data are shown as mean±s.e.m.

## 4. Discussion

The current study was designed to determine if there are relationships between expression of the stress-related neuropeptide PACAP in the extended amygdala and the social strata of group-housed C57BL/6 mice that occurs in the natural establishment of social dominance hierarchies among conspecific cagemates. Our hypothesis was that due to chronic, intermittent aggressive social interactions and naturalistic stress that animals experience over the nine weeks of group-housing, as mice self-organize into the social dominance hierarchies, there will be changes in stress-related factors that correspond with dominance rank. Social dominance rankings in the current study were based on the frequency of agonistic behaviors (e.g. aggressive and submissive behaviors) among group-housed mice following introduction into a new, unfamiliar homecage. Through these weekly perturbations in housing, mice re-establish and reveal stable dominance hierarchies that we presumed would have an underlying neurobiological signature in brain areas that subserve stress, threat detection, and adaptive responding, namely, the extended amygdala comprising the PACAP-dense regions of the BNST and CeA^48, 55, 56^. Accordingly, we found that expression of PACAP was highest in the BNSTov of the least-dominant (i.e., Submissive) mice compared to the most-dominant (i.e., Dominant) mice or mice ranked as neither most-or least-dominant (i.e., Intermediates). Dominant and Intermediate group animals showed similar levels of PACAP in the BNSTov. Interestingly, there were no differences in PACAP expression in the CeAL between any of the social dominance groups. Further, we examined if there was any impact of social dominance hierarchy on other dependent measures that are known to be influenced by stress and PACAP including body weight, CORT levels, and the acoustic startle response. Although Submissive mice tended to have lower body weight, there were no significant differences between groups. We found that while cage change and the initiation of agonistic behaviors among animals stimulated CORT in both Dominant-and Intermediate-ranked mice, CORT was significantly lower in Submissive animals compared to the other two groups. Despite these changes in PACAP and CORT, we did not observe any differences between groups in sensorimotor reactivity of the acoustic startle response.

The finding of elevated PACAP expression in the BNST of Submissive group mice — cagemates that are the recipients of the greatest amount of aggressive behavior and respond with the greatest amount of defensive behavior — is consistent with previous reports showing that repeated bouts of stress increase both mRNA transcript and protein levels of PACAP in the BNST of rodents^36, 57, 58^. Unlike chronic stress, however, the effects of a single exposure to stress on PACAP levels in the BNST are mixed, as acute restraint stress has been shown to have no effect on PACAP mRNA levels^36^, whereas exposure to a series of footshocks significantly increases the density of PACAP peptide expression^50^. Differences in the quantitative and qualitative experience of the stress is likely to account for differential effects on adaptive regulation of PACAP in the BNST. In the current study, noxious visceral stimuli (e.g., bites, anogenital sniffing, overgrooming) received by Submissive group mice would presumably be transmitted to the dorsal vagal complex (DVC) and lateral parabrachial nucleus (LPBN), where PACAP neurons reside and send projections to the BNST and CeA^46, 59–62^. Hence, adaptations within this ascending circuit as a consequence of the repeated physical stress of aggressive engagements could account for the upregulation of PACAP peptide in the BNST seen in the Submissive group mice. While the BNST is traditionally known as a node for the integration of exteroceptive and interoceptive input related to stress that can regulate autonomic, neuroendocrine and behavioral phenotypes resembling anxiety states^63–66^, recent reports indicate that it also plays a role in coordinating social behavior^67^. We cannot determine from the current study as to the functional significance of increased PACAP expression in the BNST of Submissive group mice, but speculate that it may represent a neuroadaptation that provides some survival advantage to a low-ranking animal in the social dominance hierarchy.

Along these lines, previous studies have shown that intra-BNST infusions PACAP elevates CORT levels in the blood^36^ and enhances acoustic startle reactivity^50, 57^, which could be advantageous in helping animals increase arousal, attention and escape responding^68, 69^. In contrast, other reports have shown that intra-BNST infusions of PACAP produce anorexia and loss of body weight which, at face value, would appear as maladaptive with potentially deleterious effects for the animal^70^. To better understand how our own findings of increased PACAP expression in the BNST in Submissive group mice fit in the context of these previous reports, we further examined the impact of homecage social dominance hierarchy on body weight, blood CORT levels, and acoustic startle (see below).

Our finding that PACAP expression levels in the CeA were not different between any of the social dominance groups, including Submissive group mice, aligns with a previous report showing that chronic stress (intermittent exposure to variable stressor) upregulates PACAP mRNA signal in the BNST, but not the CeA^57^. Further, others have shown that withdrawal following chronic intermittent ethanol dependence increases PACAP levels in the BNST but not the CeA^71^. Hence, there is an apparent dissociation between PACAP upregulation in the BNST versus the CeA which may be related to the qualitative nature of the type of stress experienced by animals. Despite that fact that both the BNST and CeA have been conceptualized as an interconnected neural continuum with similar afferent and efferent connections, including PACAPergic innervation, a dissociation between functional roles for these brain areas has been proposed^72^. According to this construct, the BNST plays a predominant role in sustained threat monitoring and may subserve behavioral phenotypes akin to anxiety (sustained fear) whereas the CeA is preferentially sensitive to phasic threats, mediating behaviors akin to acute fear^66, 73^. In this context, and in relation to homecage social dominance hierarchies where subordinate animals may experience chronic intermittent stress as social rank is continually challenged and reinforced (i.e. sustained threat), we speculate that experience-dependent activation of BNST circuits, including PACAPergic pathways, may predominate over CeA circuits to transform the brains of animals of the lowest social dominance rank.

We also examined the impact of homecage social dominance rank on body weight given reports that, in general, group-housed subordinate animals have lower body weight than dominant animals^5, 10, 14, 74^. While we did see a trend for Submissive group mice to have lower body weight than Dominant or Intermediate group mice, these differences were not significant. Further, we examined the relationship between BNSTov PACAP expression and body weight across all social dominance groups; there was a trend for animals with lower body weight to have higher PACAP expression in the BNSTov, but this correlation was not significant. Given that intra-BNST infusion of PACAP has been shown to induce anorexia and body weight loss^70^, we might have expected to see significantly lower body weight in Submissive group mice given our finding that these animals have increased BNSTov PACAP expression. However, a relatively high concentration of intra-BNST PACAP was required to induce significant weight loss (1.0 μg) in the prior report^70^ and it is unclear how comparable that is to the increase in endogenous PACAP in the BNSTov we observed in Submissive group mice such that it could have significantly impacted food intake and corresponding body weight in this group. Rotorod performance across all three dominance group was not significantly different indicating no impact of social dominance rank on motor coordination, as expected, based on previous reports assessing general motor performance in dominant versus subordinate animals^11, 14, 15^. We also examined the impact of social dominance hierarchy on the acoustic startle response, a sensorimotor reflex that is sensitive to repeated stress and anxiety-like states^31, 49^ and is significantly increased by intra-BNST infusion of either CRF or PACAP^50, 57, 75^. However, we did not observe any differences in acoustic startle response between any of the social dominance groups.

Assessment of blood CORT is frequently used as an assay to explore the physiological effects of social dominance as it reflects activity of the descending hypothalamic-pituitary-adrenal (HPA) axis and has traditionally been thought of as a biomarker for stress and healthy functioning of this important feedback system^76^. Published data, however, are mixed as to the relationship between CORT levels and social dominance rank as some reports indicate subordinate animals have higher levels of CORT than dominant animals, whereas others report the opposite, or no change at all^74, 77^. Differences between findings are likely related to the use of different strains, methods for dominance assessment, and stress-history of the animals^11^. In the current study we found significantly lower CORT levels in Submissive group animals compared to Dominant and Intermediate group animals in the condition where animals had just engaged in the agonistic behaviors that defined their rank (i.e., 15 min following cage change at Week-12, which is intended to serve as an activating stimulus). In comparison to a control cage of group-housed mice sacrificed without the cage change stress, Submissive group mice had a blunted CORT response compared to the other two dominance groups. A related finding has been reported in rats showing that a subgroup of subordinate-ranked mice failed to mount the appropriate surge in CORT levels following exposure to a stressor^5^. This subgroup of “non-responders” made up approximately 40% of mice ranked as subordinate, suggesting that subordination can have profound effects on the adaptive function of the neuroendocrine system responsible for an organism’s ability to respond to stressful conditions^5^. In a more recent report using an assessment to rank group-housed male mice (4/cage) as dominant or subordinate (e.g. scoring of homecage agonistic behavior) similar to ours, the mice ranked lowest in the dominance hierarchy had lower serum CORT levels than dominant mice, although the effects of acute stressor on CORT was not assessed to determine if there is a blunted response in subordinate mice^14^.

Given our observation of increased PACAP expression in the BNSTov of Submissive group mice, it is tempting to propose that there may be a causal relationship between this neuroanatomical finding and the finding of reduced CORT levels in this same subgroup. While it is known that the BNST plays a role in descending control of the HPA axis response to stress^78^, the heaviest projections from the BNST to the paraventricular nucleus of the hypothalamus (PVN; the first relay in the descending HPA axis) appear to originate from other subdivisions of the BNST, including the anterior ventral (av) and dorsomedial (dm) BNST, with only a weak projection from the BNSTov^79–84^. However, there is evidence that CRF neurons in the BNSTov do project directly to the PVN in mice and rats^85^. This projection is likely relevant because PACAPergic afferents in the BNSTov make direct contact with CRF neurons in this area^86^ and presumably influence their activity. Further, because CRF neurons in the BNST are known to also co-express GABA^87^, it is possible that PACAP in the BNSTov modulates inhibitory tone in the PVN through this same projection^64, 88^. Hence, enhanced PACAPergic innervation of the BNSTov, as seen in Submissive group mice, might affect PVN activity via enhanced CRF/GABA release and potentially serve to inhibit activity of the HPA axis to blunt CORT release in response to stress. Such a mechanism, however, would need to be reconciled with seemingly contradictory findings that 1) direct intra-BNST infusion of PACAP can significantly increase CORT levels^36^ and 2) elevated CORT in response to social defeat stress is attenuated in PACAP knockout mice^89^. In addition, it has been reported that another source of PACAP innervation of the BNST is from the PVN itself, suggesting the potential for reciprocal control of BNST-PVN feedback circuits under the control of PACAP^62^. Our current studies provide the basis for future mechanistic studies to characterize these circuits and their roles in social dominance.

Clearly, the multiple parallel and intersecting systems that underlie adaptive control of stress and may be recruited in the development of social dominance is more complex than can be addressed here. However, we believe our current findings contribute new insight into the study of social dominance and the influence of stress peptides, such as PACAP, on these networks^9, 51^. Of interest to us is how these systems may become pathologically dysregulated as a consequence of sustained threat and chronic intermittent stress like that experienced by subordinates in the context of social dominance hierarchies. Bullying in children and teenagers is one such manifestation of social dominance that has demonstrable adverse effects on mental health in subordinated youth, with greater incidence of depression, anxiety, self-harm and suicidality in victims^90, 91^. Hence, developing preclinical models with improved face and construct validity to study the impact of social dominance may lead to transformative advances in our understanding of human psychiatric conditions associated with social dominance and subordination and enable studies in research animals to better predict outcomes in humans. While we did not uncover a behavioral phenotype (e.g. changes in acoustic startle) in the current study, we did find two significant effects that were idiosyncratic to submissive animals: 1) increased expression of PACAP peptide in the BNSTov and 2) blunted CORT response following mild stress instigated by agonistic interactions. The later finding is interesting as blunted cortisol reactivity in response to stress has been associated with major depression, including in children^92, 93^. Taken together, these observations made in low dominance-ranking animals may have translational value in an effort to elucidate pathologies associated with the adverse consequences of social dominance in humans, and potentially identify better treatments for disorders that arise from these experiences.

## Acknowledgments and Disclosures

This research was supported by National Institutes of Health grant P50MH115874 (WAC and KJR). Within the past 2 years, WAC has served as a consultant for Psy Therapeutics, and has had sponsored research agreements with Cerevel Therapeutics and Delix Therapeutics that are unrelated to the current studies.

---

**Supplementary Figure 1.**
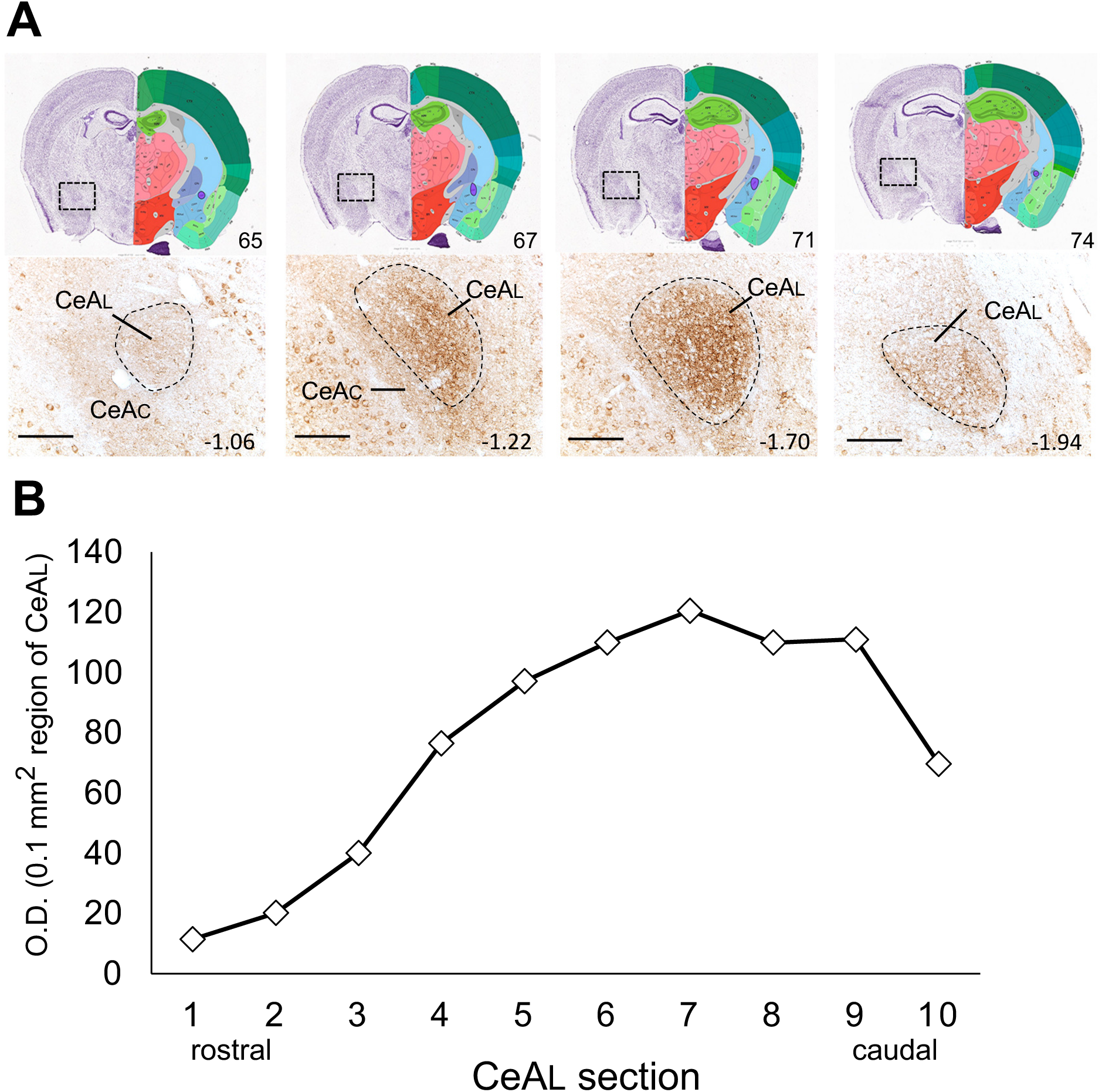
Representative coronal sections through the rostro-caudal extent of the CeA showing differential expression levels of PACAP. **(A)** *Top panel:* representative images from the Allen Mouse Brain Atlas showing the CeA at different rostro-caudal (left to right) levels; numbers in lower right corner of each image refer to the plate image from the Allen Brain Atlas mouse.brain-map.org and atlas.brain-map.org. ^95, 96^. Boxed area on left side (Nissl stained) of image represents magnified section in lower panels of **A** showing PACAP expression (brown immunolabel) and approximate boundaries of CeAL at different rostro-caudal levels. Numbers in lower right corner of each image refer to the approximate brain level shown in millimeters posterior to bregma from the mouse brain stereotaxic atlas of Paxinos and Franklin (2019)^97^ for reference. **(B).** Optical density (O.D.) measurements of 0.1 mm^2^ regions through the CeAL from serial (every third) sections through the CeA. As illustrated in the line graph, PACAP expression was strongest in caudal sections of the CeA. Scale bars = 100 μm.

